# Repeated birth injuries lead to pelvic floor muscle dysfunction and impairment in regeneration

**DOI:** 10.1101/2022.05.02.490259

**Authors:** Pamela Duran, Emma Zelus, Saya French, Lindsey Burnett, Karen L. Christman, Marianna Alperin

## Abstract

**Objectives:** Childbirth is a key risk factor for pelvic floor muscle (PFM) injury and dysfunction, and subsequent pelvic floor disorders (PFDs). Multiparity further exacerbates these risks. Using the pre-clinical rat model of simulated birth injury (SBI), we previously identified that an SBI leads to PFM atrophy and fibrosis. We hypothesized that multiple SBIs further overwhelm PFM regenerative capacity, leading to functionally relevant pathological alterations long-term.

**Study Design:** Rats underwent SBI and were allowed to recover for 8 weeks to undergo another SBI. Animals were sacrificed at acute, subacute, and long-term time points post-second injury (N=3-6/time point), and pubocaudalis (PCa) was harvested to assess *ex vivo* muscle function, histomorphological properties and gene expression.

**Results:** Acutely following the 1^st^ SBI, PCa force was decreased relative to controls. At 4 weeks, PCa force was recovered and remained unchanged at 8 weeks. Similarly, lower PCa force was observed immediately after repeated SBI. In contrast to functional recovery after 1^st^ SBI, PCa force remained lower at 4 weeks post-2^nd^ SBI and continued to be decreased even after 12 weeks after repeated injury. Fiber size was smaller at the long-term time points after 2^nd^ SBI compared to controls and single SBI groups. As opposed to the resolution of centralized nuclei at 8 weeks post-1^st^ SBI, regenerating myofibers persisted even at 12 weeks post-2^nd^ SBI. In contrast to the peak of collagen content at 4 weeks post-1^st^ SBI, this parameter raised progressively over 12 weeks after repeated SBIs. Prolonged inflammatory response, impairment in muscle anabolism, and sustained expression of ECM remodeling genes were observed after repeated SBIs.

**Conclusions:** Repeated birth injuries delay PFM regeneration and impair function in the pre-clinical rat model.

## Introduction

Vaginal delivery has been consistently identified as a key risk factor for pelvic floor muscle (PFM) injury and the associated pelvic floor disorders (PFDs)^1^. In addition, multiparity further increases this risk, especially for the development of pelvic organ prolapse (POP)^2-5^. Despite the established epidemiological risk factors, the mechanisms leading to PFM dysfunction in primiparous and multiparous women remain elusive.

We previously discovered that despite an 8-week recovery period following vaginal distention, pubocaudalis (PCa)—analogous to human pubovisceralis^6^—exhibits atrophic and fibrotic pathological alterations in the rat pre-clinical model^7^. The above morphological aberrations are associated with sustained inflammatory response, impairment in muscle anabolism, and persistent extracellular matrix remodeling^7^. To investigate whether these long-term pathological alterations impact PFM function, the first objective of the current study was to assess how simulated birth injury (SBI) affects mechanical muscle properties.

Studies investigating the effect of repeat injuries on the limb muscle recovery demonstrate profound muscle atrophy associated with loss of regenerative capacity^8, 9^. To our knowledge, there are no published studies examining the impact of repeat large mechanical strains on the female PFMs. In a previous study, focused on the effect of multiple vaginal distentions on external urethral sphincter recovery, the investigators observed a profound decrease in leak point pressure after repeated SBIs in the rat model^10^. However, it is not clear whether active regeneration of the striated urethral sphincter was completed at the time of the repeated injury in the above study. Several investigations have indicated that muscle response is dramatically affected when a second injury is performed during active muscle regeneration^8, 11^. Thus, the second objective of the current study was to test the following hypotheses: 1) repeated SBI, imposed on the PFMs after completion of an active recovery period following the first SBI, further overwhelms PFM regenerative capacity, leading to more substantial pathological alterations than a single birth injury and 2) pathological alterations in the intrinsic PFM components incited by repeated SBIs result in PFM dysfunction, a known risk factor for clinically relevant PFDs observed in multiparous women.

In this study, we found that repeated SBIs lead to increased pathological alterations and decreased active force production in relation to a single injury. In addition, an inflammatory response, impairment in muscle anabolism, upregulation of genes associated with muscle catabolism, and sustained ECM remodeling were observed after repeated birth injuries.

## Materials and methods

### Study design

The objectives of the current study were to: 1) assess PFM function after a single SBI to correlate with previously identified pathological alterations, 2) determine the impact of repeated birth injuries on PCa function, histological alterations, and gene expression profile. A 2^nd^ SBI was performed 8 weeks after the 1^st^ SBI, since muscle regeneration was completed at this time point^7^. The sample size, indicated in the figure legend, was calculated based on preliminary experiments. No outliers were excluded from the study. For the quantitative analyses, the investigators were blinded to the group identity.

### Pelvic floor muscle mechanics

#### Active mechanical testing

Animals were anesthetized with 3% isoflurane @ 2 L/min; a midline incision was made at the level of the symphysis pubis, which was disarticulated, and PCa was identified and harvested for *ex vivo* mechanical testing. The PCa portion of the rat levator ani was again the focus of these experiments, as this muscle’s stretch ratios are similar to pubococcygeus, which experiences the largest strains during human parturition^12, 13^. Immediately after dissection, animals were euthanized by CO_2_ inhalation, followed by bilateral thoracotomy. The muscle was placed in an organ bath (805-A, Aurora Scientific, Aurora, Ontario) with aerated (95% O_2_, 5% CO_2_) mammalian Ringer’s buffer (137 mN NaCl, 5mM KCl, 24 mM NaHCO_3_, 1mM NaH_2_PO_4_, 11mM C_6_H_12_O_6_, 1mM MgSO_4_, 2mM CaCl_2_, pH 7.4) and platinum-based electrodes (Fig. 4.1B). Using 4.0 silk sutures, attached to the proximal enthesis and distal tendon, PCa was attached to the fixed post on one side and the lever arm of the force transducer (300C: Dual-Mode, 1.5 N, Aurora Scientific, Aurora, Ontario) on the other side (Fig. 1A-B). For all mechanical testing, electrical pulses were delivered to the platinum-based electrodes using PowerLab (4/35 Power Lab; ADI Instruments, Sydney, Australia) and an electrical stimulator (701C, Aurora Scientific, Aurora, Ontario). Force output was recorded from the force transducer with LabChart (v8.1.3; ADInstruments, Sydney, Australia).

**Figure 1.**
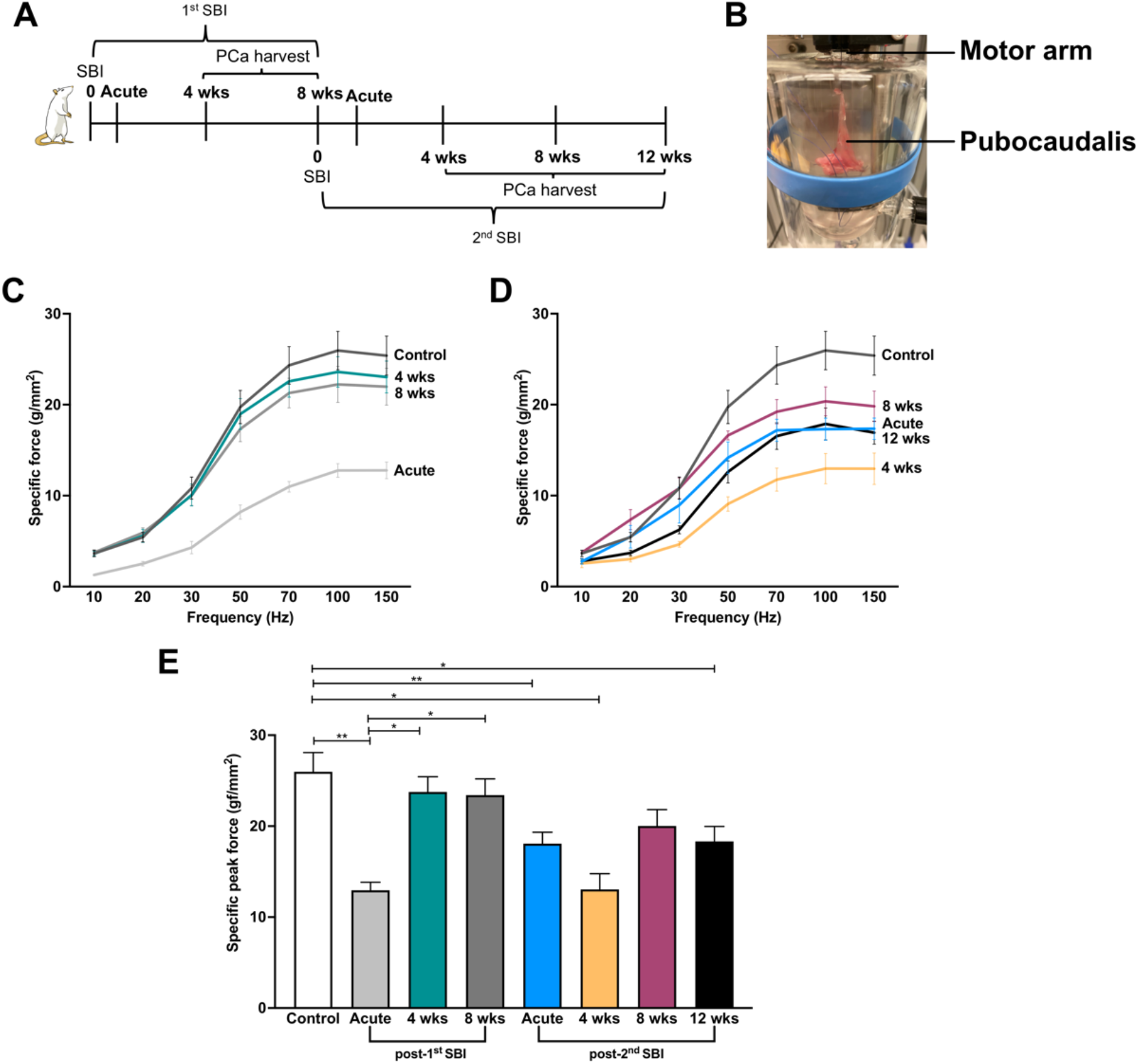
Pelvic floor muscle active function after single and repeated simulated birth injuries. (A) Timeline of experimental design. (B) Example of pubocaudalis (PCa) in organ bath attached to the force transducer by the tendon at the insertion and fixed to the arm with the pubic bone. Force-frequency curves at various timepoints after a single (C) or repeated (D) simulated birth injury and specific peak force (E). N=3-7/group. *P*-values derived from two-way ANOVA followed by pairwise comparisons with Sidak’s range test. *p<0.05, **p<0.01, ***p<0.001, mean±SEM.

To determine muscle length corresponding to the greatest force generation, known as the optimal muscle length (L_o_), PCa was gradually stretched to achieve 0.05 V difference per increment, starting from the voltage at slack muscle length. PCa was stimulated with isolated 1 Hz electrical pulses at each muscle length until maximal force was determined^14^. Once L_o_ was identified, PCa was subjected to a range of stimulations to establish the force-frequency relationship in uninjured and injured animals. Electrical stimuli (0.2 ms, 1 A, 1200 ms trains) were applied with successive frequency increases (10, 20, 30, 50, 70, 100, 150 Hz). Each stimulus was interspersed by a two-minute rest interval. Maximum force at each frequency was normalized by physiological cross-sectional area, which was calculated using validated methods^14, 15^. Briefly, muscle mass was divided by the product of fiber length (L_f_) and muscle density (1.06 mg/mm^3^)^15^. L_f_ was obtained by multiplying L_o_ by the previously determined PCa muscle length to fiber length ratio of 1.68^6^. After normalization, force-frequency curves and peak force were obtained.

#### Passive mechanical testing

After active mechanics experiments, the PCa was placed in a relaxing solution with leupeptin, a protease inhibitor, and stored at -20°C for at most 2 weeks^16^. The PFM was then micro-dissected to obtain small muscle fascicles—3 fascicles/muscle were analyzed. Laser diffraction was used to measure the optimal sarcomere length (L_s_) and calipers were used to measure the fascicle length at that L_s_. Each fascicle was then attached to the force transducer and the fixed post at the previously measured fascicle length. Tissue was evaluated by stress-relaxation tests, where the muscle was stretched at a rate of 2 L_s_/s in 10% increments until failure or to 60%, with 3-min relaxation period between increments^17^. Force values were divided by cross-sectional area— assuming cylindrical shape^16^—to obtain stress-strain curves. Sample stiffness was calculated by measuring the slope of the stress-strain curve at 20, 30, 40 and 50% stretch.

### Immunohistochemical analyses

PCa was harvested, embedded in optimal cutting temperature media, and snap-frozen in isopentane chilled in liquid nitrogen. Samples were cryosectioned at 10 μm and prepared for immunohistochemical analyses. To assess fiber area and centralized nuclei, slides were fixed in 4% PFA, rinsed three times in 1x phosphate buffer saline (PBS) and incubated in blocking buffer for 1 hour (10% normal goat serum (Gemini Bio-products, Sacramento, CA) in PBS). Slides were then incubated overnight with anti-laminin antibody (Abcam, Cambridge, MA; 1:200), rinsed in PBS and incubated in Alexa Fluor 488 conjugated secondary antibody (Invitrogen, Carlsbad, CA; 1:500) for 30 minutes. After rinsing in PBS, nuclei were counterstained with Hoechst 33342 (1: 10000). Slides were mounted with Fluoromount (Sigma-Aldrich, St. Louis, MO) and scanned at 20X on the Ariol DM6000 B microscope (Leica, Wetzlar, Germany). Fiber area was quantified using a custom macro in ImageJ (NIH, Bethesda, MD) for a total of ∼10,000 fibers per sample obtained from randomly selected 4-8 tissue sections. Centralized nuclei per total number of fibers were counted manually.

For the quantification of the intramuscular collagen content, slides were fixed in cold acetone, rinsed three times in 1xPBS and incubated in blocking buffer (10% normal goat serum (Gemini Bio-products, Sacramento, CA) + 0.3% Triton X-100 (Sigma-Aldrich, St. Louis, MO) + 1% bovine serum album (Sigma-Aldrich, St. Louis, MO) in PBS)) for 1 hour. Slides were then incubated overnight with collagen I primary antibody (Invitrogen, Carlsbad, CA, PA595137, 1:200), rinsed in PBS, and incubated with Alexa Fluor 488 conjugated secondary antibody (Invitrogen, Carlsbad, California; 1:500) for 2 hours. Slides were then rinsed in PBS and nuclei were counterstained with Hoechst 33342 (1:10000). Slides scanned at 20X (Ariol DM6000 B microscope, Leica, Wetzlar, Germany) were processed for quantification using freehand selection tool in ImageJ (NIH, Bethesda, MD). Collagen content was expressed as area fraction occupied by collagen staining.

### Gene expression profile after repeated birth injuries

PCa was harvested at several relevant time points to assess potential causes behind the identified structural alterations and muscle dysfunction. RNA isolation, and Nanotring nCounter analysis were done as previously described^7^.

### Statistical Analysis

For the primary mechanical outcome of interest, the specific peak force, sample size calculation (G*power software) yielded 7 animals/time point based on Cohen’s d effect size of 1.7 obtained from the preliminary experiments, to achieve 80% power at a significance level of 0.05. For histological outcomes, Cohen’s d effect size of 5.2, obtained from our previous study comparing morphological outcomes between animals subjected to a single SBI and unperturbed controls^7^, was used to determine sample size. Power calculation yielded 3 animals/time point to achieve 80% power at a significance level of 0.05. Data with a parametric distribution were compared with a two-way ANOVA followed by Tukey’s or Sidak’s post hoc pairwise comparisons, when indicated. Fiber area, which followed a non-parametric distribution, was analyzed with Krukal-Wallis test followed by Dunn’s pairwise comparisons. Data were analyzed using GraphPad Prism v8.0, San Diego, CA. Data processing for gene expression was performed using the NanoStringDiff package in R, with a significance at a p value<0.05 and a fold change cutoff of 1±0.25^18^.

## Results

### The impact of single and repeat simulated birth injuries on the pelvic floor muscle function

To determine whether myofiber atrophy and fibrotic degeneration of the PFMs consequent to a single SBI are functionally relevant, we performed *ex vivo* active mechanical testing (Fig. 1A-B). Immediately following the 1^st^ SBI, PCa demonstrated a decrease in the force-frequency curve (Fig. 1C), with significant reduction in the specific peak force compared to uninjured controls (P<0.01, Fig. 1E). At 4 weeks post-1^st^ SBI, the pattern of the force-frequency curve (Fig. 1C) and the specific peak force (Fig. 1E) were similar to controls despite ongoing muscle regeneration (P=0.99) (Fig. 1C, E). These mechanical parameters remained unchanged at 8 weeks post SBI, when muscle regeneration was completed, compared to the 4-week time point (P>0.99, Fig. 1C, E). Taken together, these data indicate that despite atrophy and fibrosis—which are known to affect active muscle mechanical properties^19, 20^—of the PCa in response to a single SBI, muscle force production was not affected long-term.

Given the above and motivated by the epidemiological studies that identify increased incidence of pelvic floor dysfunction and the associated PFDs in multiparous women compared to women with history of only one vaginal delivery^2^, we went on to assess PFM function after repeat birth injuries. Immediately after repeat SBI, the force-frequency curve demonstrated decreased force production compared to uninjured controls, with a reduction in peak force (P<0.01, Fig. 1D, E). In contrast to the functional muscle recovery observed 4 weeks after a single SBI, PCa force was significantly decreased at 4 weeks post-2^nd^ SBI relative to uninjured controls (Fig. 1D, E). A reduction in muscle force was identified even at 12 weeks after 2^nd^ SBI (P<0.01, Fig. 1D, E). These results indicate that repeated birth injuries deleteriously affect PFM contractile function.

Besides assessing active mechanical properties, investigation of passive parameters is essential as a pathological increase in collagen (i.e., fibrosis) can affect muscle stiffness—which is known to affect muscle force transmission^19^. To assess this, we performed passive mechanical testing at the fascicle level (Fig. 2A, B). Interestingly, although active muscle function was not affected after a single SBI, at 8 weeks post-1^st^ SBI we observed an upward shift of the stress-strain curve compared to uninjured controls (Fig. 2C) together with an increase in stiffness at low strains (Fig. 2D). This pattern was also observed at 12 weeks post-2^nd^ SBI (Fig. 2C, D), indicating that a single or repeated birth injury affect muscle passive mechanical properties.

**Figure 2.**
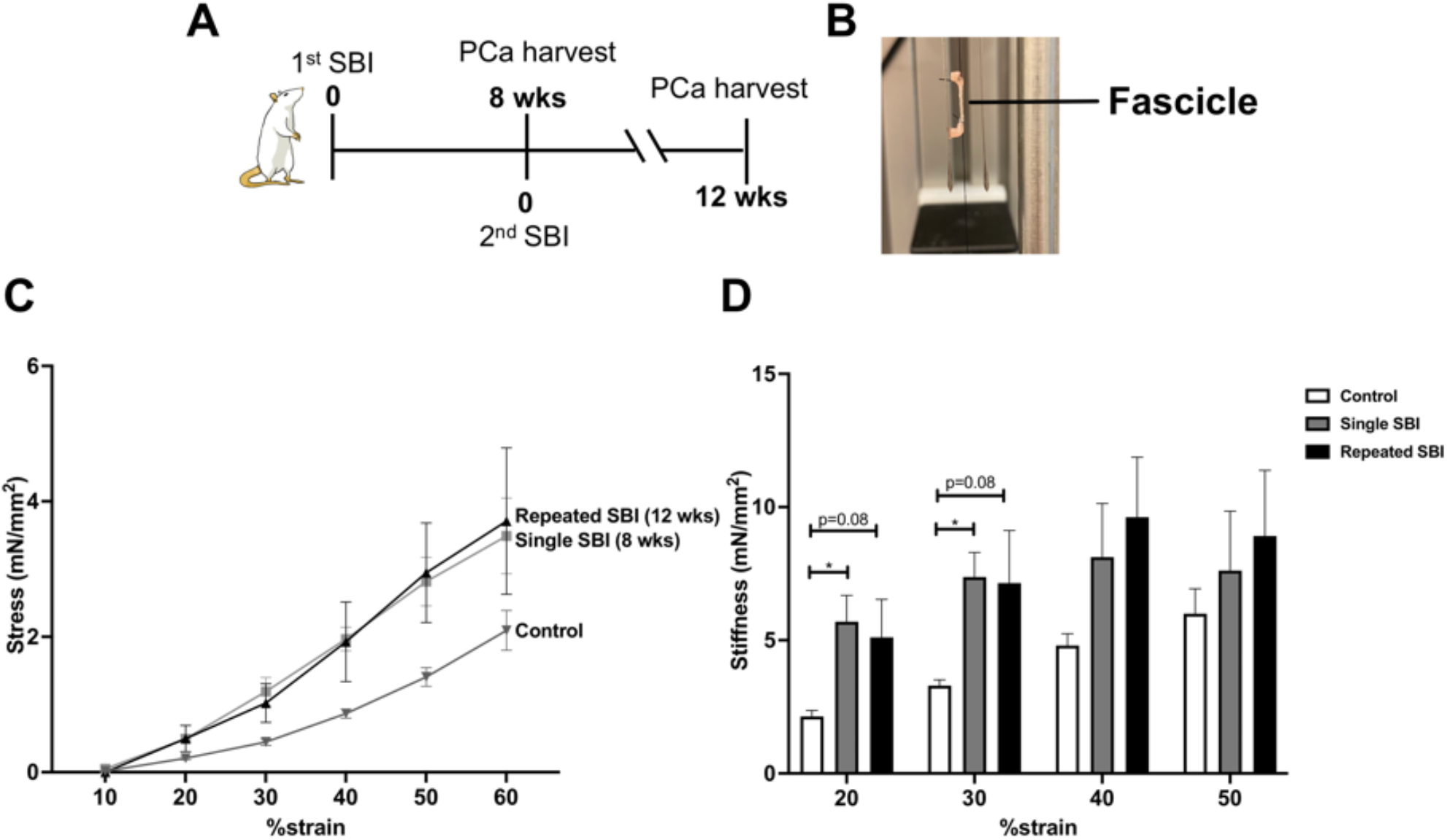
Passive mechanical properties of the pelvic floor muscle after a single or repeated simulated birth injuries. (A) Timeline of experimental design. (B) Example of fascicle attached to motor arm for mechanical testing. Stress-strain curves (C) and stiffness (D) were obtained after a single or repeated birth injury. N=3/group. P-values derived from one-way ANOVA followed by pairwise comparisons with Dunnett’s range test. mean±SEM.

### Repeated birth injuries negatively impact pelvic floor muscle regeneration

As the first step in determining potential causes of the observed PFM dysfunction after repeated SBIs, we went on to assess the impact of repeated birth injuries on PFM histomorphometric properties (Fig. 3A) and to compare to the findings in animals subjected to a single SBI^7^.

**Figure 3.**
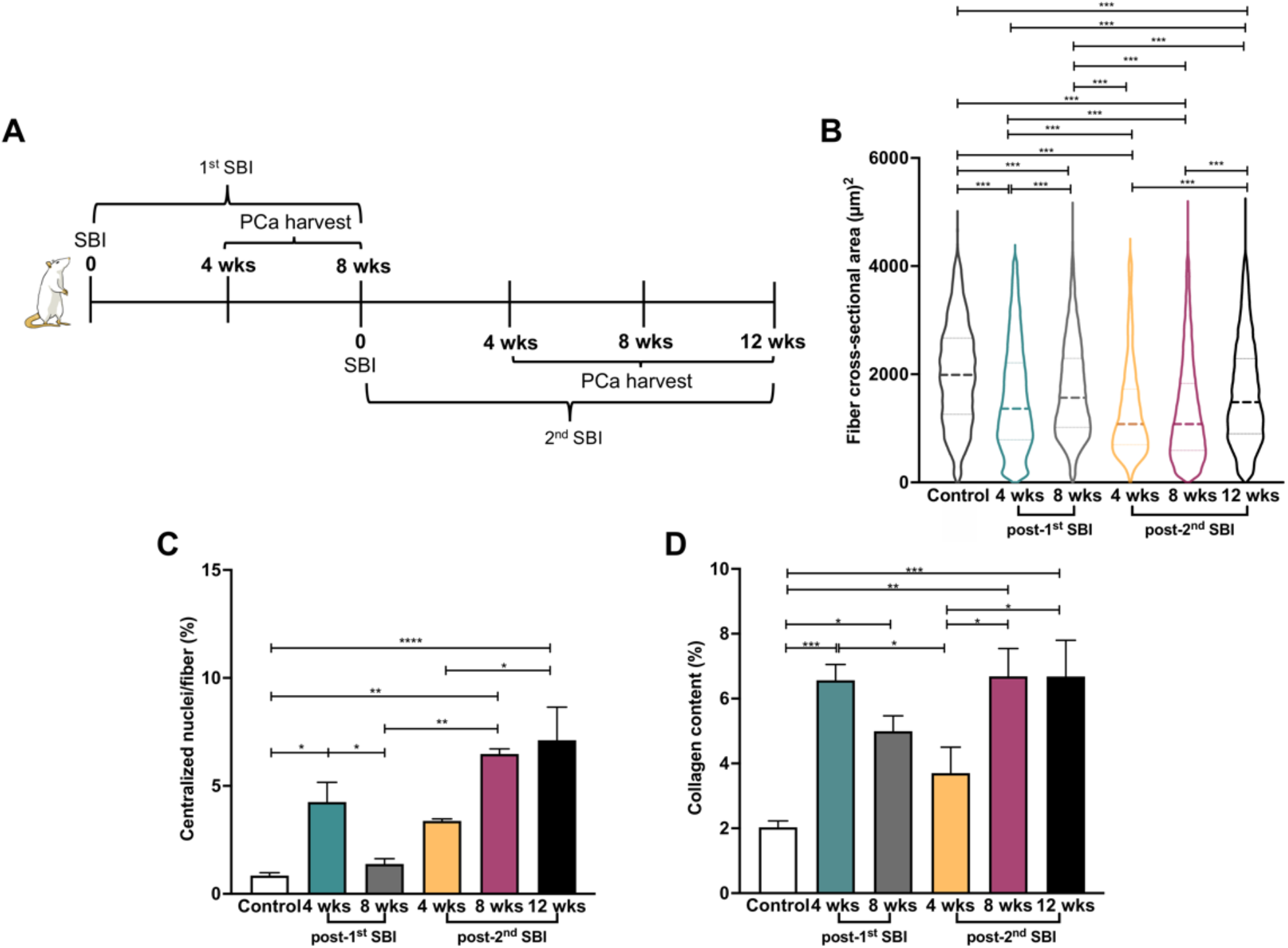
Repeated simulated birth injuries (SBI) lead to an impairment in pelvic floor muscle regeneration. (A) Timeline of experimental design. Quantification of fiber cross-sectional area (B), centralized nuclei (E) and collagen content (F) after a single or repeated birth injury. N=3-6/group. *P*-values derived from two-way ANOVA followed by pairwise comparisons with Sidak’s range test. *p<0.05, **p<0.01, ***p<0.001, ****p<0.0001, mean±SEM.

PCa fiber size decreased at 4 weeks post-2^nd^ SBI compared to controls and to animals allowed to recover for 4- or 8-weeks post single SBI (P<0.0001, Fig. 3B). Fiber area distribution did not change between 4 and 8 weeks following the 2^nd^ injury (P>0.9, Fig. 3B). This was in contrast to the findings following a single SBI, after which PCa fiber size increased between 4 and 8 weeks following the injury. We further analyzed fiber area at 12 weeks after repeated injury as we also observed a reduction in force production at this time point. While fiber size was increased at 12 weeks in relation to 8 weeks post-2^nd^ SBI (P<0.0001), fiber area was still decreased compared to uninjured controls (P<0.0001, Fig. 3B).

To assess muscle regeneration, we quantified the proportion of fibers with centralized nuclei—an index of regeneration^21^. The number of fibers with centralized nuclei post-2^nd^ SBI was not different at 4 weeks following the injury compared to controls (P=0.5, Fig. 3C). At 8 weeks after the 2^nd^ injury, the number of regenerating myofibers was substantially increased compared to uninjured controls (P<0.01, Fig. 3C). As opposed to the impact of a single SBI, after which the percent of regenerating myofibers returned to baseline by 8 weeks post injury, the proportion of centralized nuclei remained high at this time point after the 2^nd^ SBI (P<0.01, Fig. 3C). Since these results indicated ongoing muscle regeneration at this time point, we analyzed the proportion of regenerated myofibers at 12 weeks post-repeated injury. Surprisingly, even at this long-term time point, the percent of centralized nuclei was still increased in relation to controls (P<0.0001), without differences compared to the 8-weeks period (P=0.8, Fig. 3C). All together, these findings suggest an impairment in muscle regeneration after repeated birth injuries.

We also assessed changes in the intramuscular collagen content in response to multiple SBIs, as pathological increase in collagen I, the main constituent of the intramuscular extracellular matrix, can negatively affect muscle active and passive mechanical properties^22^. At 4 weeks post-2^nd^ SBI, PCa collagen content was unchanged relative to controls (P=0.0.5) and lower than that observed at 4 weeks post-1^st^ SBI (P<0.05, Fig. 3D). Eight weeks after repeated SBI, intramuscular collagen content was significantly increased relative to controls (P<0.01) and compared to the 4-week time point (P<0.05), without differences in relation to the 12-week time point (P>0.99, Fig. 3D). These results demonstrate that fibrotic degeneration of the PFMs subjected to repeated large strains occurs later than following a single vaginal distention.

### Prolonged inflammatory response of the pelvic floor muscles after repeated birth injuries

To investigate potential mechanisms behind the profound reduction in fiber size, impairment in regeneration and muscle fibrosis after repeated injuries, we then performed a gene expression study capitalizing on a customized Nanostring panel previously utilized^7^. PCa was harvested at acute, subacute and long-term time points as with a single SBI to analyze pathways essential for muscle regeneration after an injury^23-25^. We first assessed patterns across the injury time points through a Principal component (PC) analysis (Fig. 4A). PC1 that corresponded to the highest variability, indicated the time course of return to the uninjured state after a 2^nd^ SBI.

**Figure 4.**
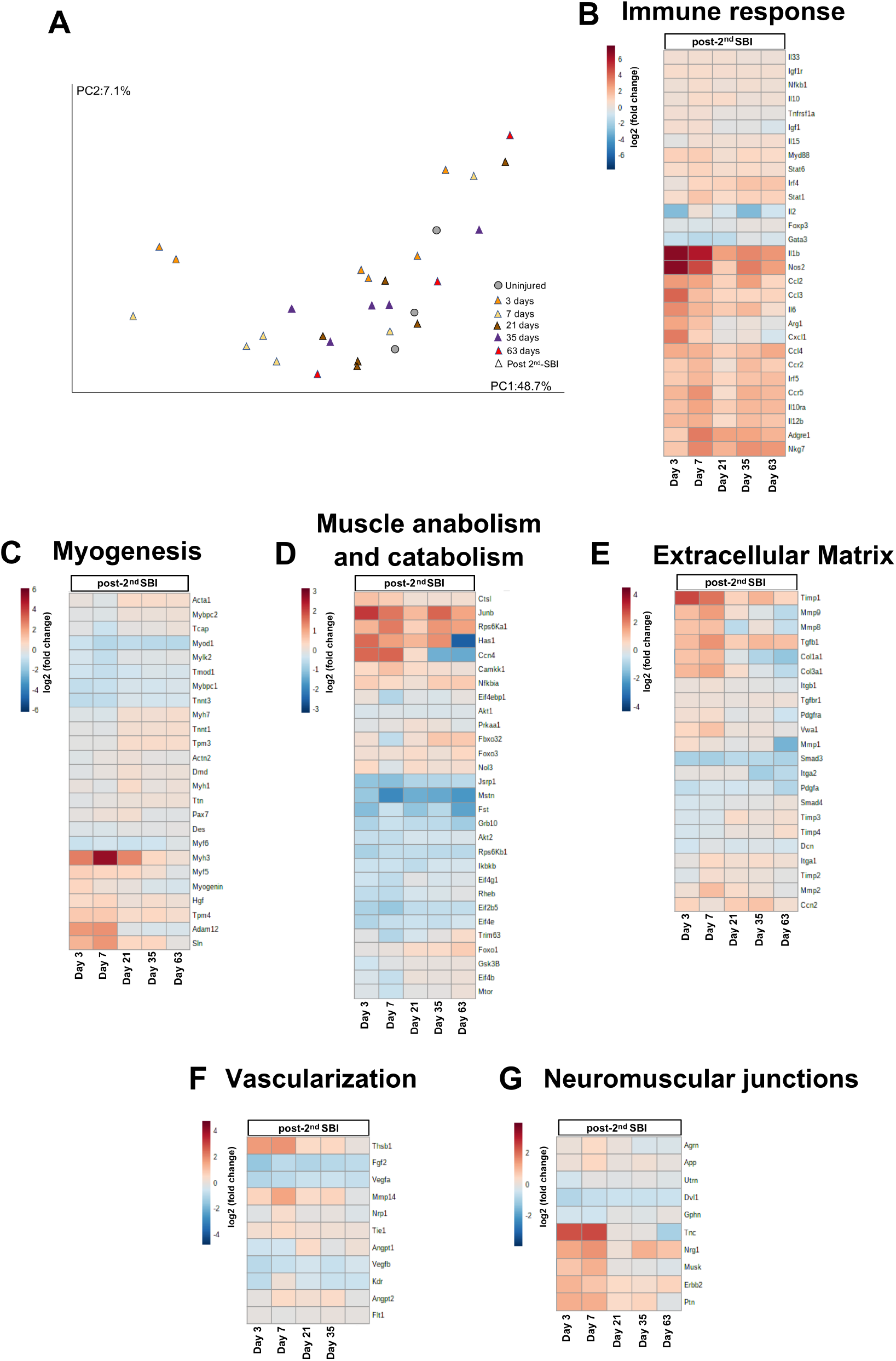
Repeated simulated birth injuries (SBI) lead to prolonged inflammatory response, sustained expression of ECM remodeling genes, impairment in muscle anabolism, and upregulation of muscle catabolism related genes. (A) Principal component analysis that includes gene expression in uninjured controls and at several time points after repeated SBI. (B-G) Unsupervised clustered heat maps of fold changes at each time point in relation to uninjured controls. Pathways include immune response (B), myogenesis (C), muscle anabolism and catabolism (D), extracellular matrix (E), vascularization (F), and neuromuscular junctions (G). N=3/6 per group.

We then investigated the longitudinal patterns in gene expression with respect to the uninjured state by analizing the differentially expressed genes (Tables S1-S6). Figure S1 indicates the gene expression profile across the replicates per timepoint post-2^nd^ SBI. Unsupervised clustered heat maps of the fold change in relation to the uninjured controls are visualized in Figure 4B-G. For the immune response category (Fig. 4B), the majority of inflammatory related genes clustered together. As with a single injury^7^, at the acute time points after 2^nd^ SBI, increased expression of pro-inflammatory genes was identified together with upregulation of some pro-regenerative genes. Prolonged expression of inflammatory genes (Il1b, Irf5, Stat1) was observed at the longer-term time point (63 days). For myogenesis pathway (Fig. 4C), the gene expression pattern was similar after a repeated injury as with a single one^7^—genes related to muscle structure were downregulated at the acute time points, with upregulation of muscle transcription factors and mediators. Notably, Myf5 and Myh3, which are involved in the regeneration of myofibers, were upregulated until day 21 post-2^nd^ SBI. For muscle anabolism and catabolism category (Fig. 4D), following the previous pattern observed after a first injury^7^, repeated injuries led to decreased expression of genes associated with muscle anabolism even at the long-term time points (Eif2b5, Rps6kb1). Uniquely, upregulation of genes associated with muscle catabolism (Foxo1, Foxo3) were identified at the longer time point. In addition, genes associated with extracellular matrix (ECM) remodeling (Mmp2, Mmp9, Tgfb1, Timp1) were observed at the acute time points post-2^nd^ SBI (Fig. 4E). From these genes, the ones that promote collagen synthesis (Tgfb1, Timp1) were still upregulated at the longer time point. Vascularization-related genes were mostly downregulated following the repeated injury (Fig. 4F). At last, genes related to the development and maturation of neuromuscular junctions were upregulated within one week after 2^nd^ SBI. All together the results suggest that repeated birth injuries lead to a prolonged inflammatory response, impairment in muscle anabolism, upregulation of genes associated with catabolism, and sustained expression of ECM remodeling genes.

## Discussion

In the current study, we first wanted to assess if a single SBI impacts PCa force production since we have previously identified that in the rat model, validated for the studies of the human PFMs^6, 12^, SBI leads to muscle atrophy and fibrosis. Despite these pathological alterations, PCa long-term active force generation recovered after a single SBI. However, passive mechanical properties were affected, as indicated by an increased stiffness. Published clinical studies demonstrate progressive worsening of PFM function after each subsequent vaginal delivery^26^. Thus, we assessed the effect of repeated birth injuries on PFM force production to determine if the rat model recapitulates this phenotype. In contrast to a single birth injury, repeated SBIs resulted in a 30% decrease in PFM contractile force production despite a 12-week recovery period, together with an increased passive stress. We next investigated changes in the morphometric properties after repeated birth injuries to begin to understand the mechanisms accounting for the observed decrease in PFM function. Our results demonstrate that relative to a single SBI, repeated birth injuries at the time of an atrophic and fibrotic PFM, lead to an impairment in muscle regeneration, reduced fiber size, increased collagen content, all which can negatively affect muscle function^20^.We then assessed changes in gene expression across physiological time points to further determine possible causes for the observed histological alterations. Repeated birth injuries lead to a prolonged inflammatory response, upregulation of genes associated with catabolism, impairment in muscle anabolism and sustained expression of ECM remodeling genes.

In our study, the reduced fiber size and increase in collagen content coincide with the decrease in active muscle force production and trending increase in passive stiffness at 12 weeks post-2^nd^ SBI. In our rat model, a substantial 150% increase in collagen content after the single injury was enough to increase stiffness. It is well established in the limb muscles that muscle force production has a direct relationship to myofiber cross-sectional area^20^, however the exact extent of alterations in fiber size that impact this functional parameter remains undetermined. We observed that a 15% decrease in fiber size at 8 weeks after single SBI was not enough to influence muscle active force production, while a 25% reduction after repeated birth injuries lead to decrease active force. Taken together, these data indicate a “dose-dependent” relationship between the number of birth injuries, the extent of the pathological alterations of the PFM morphometric properties, and the impact on active muscle function. The above provides a putative mechanistic link between multiparity and progressive worsening of the PFM function identified clinically^26^.

We performed a gene expression study to identify potential causes behind the histological alterations observed after 2^nd^ SBI. With respect to a single SBI^7^, prolonged inflammatory response was observed after repeated injuries, which could also indicate such “dose-dependent” relationship between the number of birth injuries, and the presence of the altered immune response. An inflammatory response negatively influences muscle anabolism, specifically the AkT/mTOR signaling, and also enhances muscle catabolism^25^. In our study, the persistent inflammatory response observed after repeated injuries could be associated with the profound reduction in fiber size, through impairment in muscle anabolism and upregulation of catabolism. In addition, a small fiber area could also be related to the observed increase in centralized nuclei. The genes associated with myogenesis pathway were upregulated at the expected time course, elucidating the investigation of other mechanisms behind the substantial increase in centralized nuclei identified at the long-term time points.

Due to significant technical constraints associated with dissecting the pubocaudalis branch of the levator ani nerve, we relied on the *ex vivo* mechanical testing. While the results could be potentially divergent from the *in vivo* mechanics, our protocol allowed us to uncouple potential neurogenic influence on the observed changes in muscle force and focus exclusively on the independent myogenic contributions to the observed phenotype.

In conclusion, our findings indicate that repeated birth injuries lead to increased pathological alterations, and prolonged inflammatory response compared to a single birth injury. The resultant impairment of PFM force production provides potential mechanistic link behind the additive risk of multiparity for PFM dysfunction.

