## Supplementary Materials for "Repeated birth injuries lead to pelvic floor muscle dysfunction and impairment in regeneration"

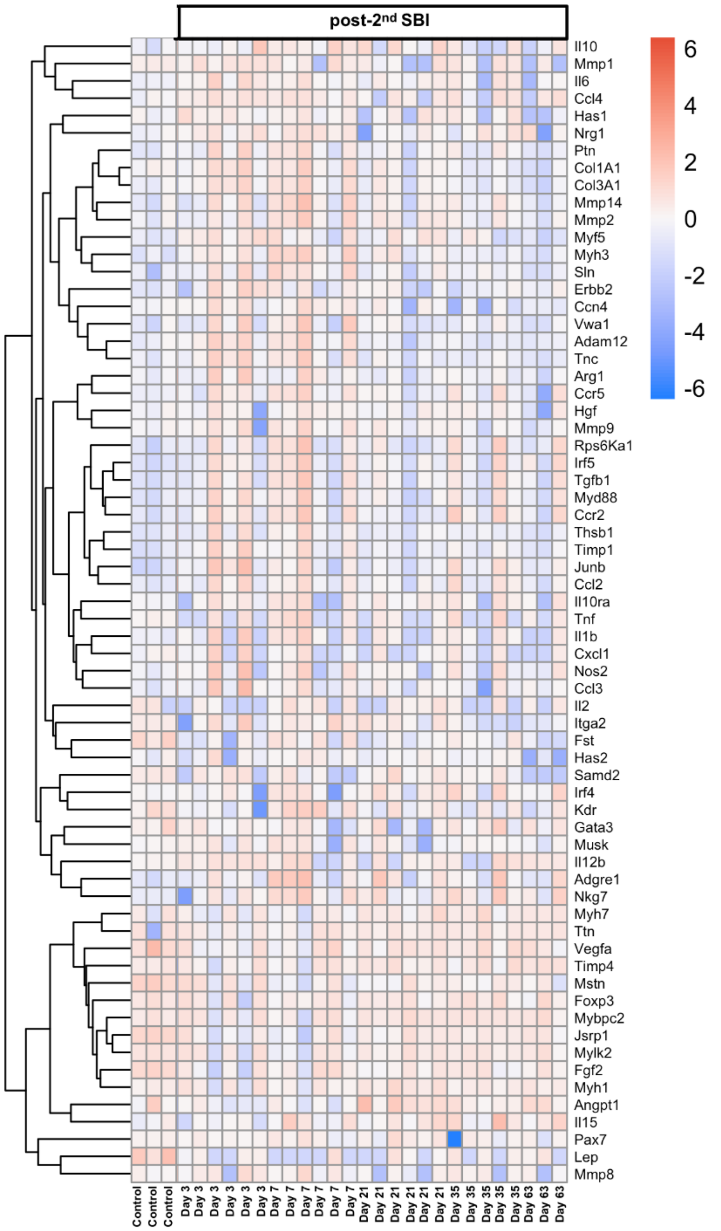


**Figure S1 Gene expression profile across the replicates per time point after repeated simulated birth injury.**

Heat map of genes with the highest variation in the dataset. N=3/6 per group.

**Table S1. Immune response related genes throughout the different time points after repeated simulated birth injuries.**

P and q values are indicated for significantly differentially expressed genes compared to uninjured group. FC= Log2FC

| | Days after 2nd Simulated Birth Injury | | | | | | | | | | | | | | |
|---|---|---|---|---|---|---|---|---|---|---|---|---|---|---|---|
| Gene | 3 | | | 7 | | | 21 | | | 35 | | | 63 | | |
| | FC | p | q | FC | p | q | FC | p | q | FC | p | q | FC | p | q |
| Adgre1 | 1.31553789 | | | 3.55627507 | 0 | 0 | 2.61525684 | 0.01 | 0.08 | 2.529661 | 0.01 | 0.08 | 2.07202034 | | |
| Arg1 | 2.47717966 | 0 | 0.02 | 1.64061668 | | | -0.1775432 | | | 0.13685368 | | | -0.2287393 | | |
| Ccl2 | 2.80655752 | 0 | 0.01 | 2.46072917 | 0.006 | 0.01 | 1.11463903 | | | 2.68913478 | 0.003 | 0.03 | 1.00945403 | | |
| Ccl3 | 4.11776569 | 0 | 0 | 1.76777647 | | | 1.01452302 | | | 1.01265361 | | | 1.03009956 | | |
| Ccl4 | 2.3352731 | 0 | 0 | 2.13016896 | 0 | 0 | 1.15190687 | | | 1.68022553 | | | 2.34035449 | 0.005 | 0.05 |
| Ccr2 | 1.34081704 | 0.02 | 0.06 | 1.90713463 | 0 | 0.01 | 0.31443285 | | | 1.80931201 | 0.004 | 0.04 | 1.27484094 | | |
| Ccr5 | 1.98280117 | 0.04 | 0.1 | 3.02550858 | 0.004 | 0.01 | 0.62804597 | | | 2.16814896 | 0.03 | 0.1 | 1.85722485 | | |
| Cxcl1 | 3.57829369 | 0.01 | 0.03 | 1.10774696 | | | -0.0616809 | | | 0.22605741 | | | -0.6684862 | | |
| Foxp3 | -0.3917188 | | | -0.5248428 | | | -0.3137782 | | | -0.041802 | | | -0.0352335 | | |
| Gata3 | -1.1133997 | | | -1.4412141 | 0.03 | 0.05 | -1.3379966 | | | -0.3238518 | | | -0.6341679 | | |
| Igf1 | 0.48787423 | | | 0.50704644 | | | -0.2652431 | | | -0.2237509 | | | -0.5481125 | | |
| Igf1r | 0.65565008 | 0 | 0 | 0.61258357 | 0.002 | 0.01 | 0.50389617 | 0.01 | 0.09 | 0.43836134 | 0.04 | 0.1 | 0.45572202 | | |
| Il10 | 0.24608613 | | | 0.68189705 | 0.01 | 0.02 | 0.70288497 | 0.01 | 0.09 | 0.18316467 | | | 0.33546007 | | |
| Il10ra | 1.69596356 | | | 2.4842666 | 0.01 | 0.03 | 1.31647514 | | | 1.96860543 | | | 1.90944274 | | |
| Il12b | 1.89969165 | 0.01 | 0.04 | 2.2159074 | 0 | 0 | 0.74877339 | | | 1.75069959 | 0.04 | 0.1 | 1.523696 | | |
| Il15 | -0.198507 | | | 0.4068818 | | | 0.30165052 | | | 0.39873005 | | | 0.5367961 | | |
| Il1b | 6.88124101 | 6.88E-14 | 2.39E-12 | 5.95816588 | 1.77E-15 | 8.23E-14 | 2.68458831 | 2.26E-14 | 1.04E-12 | 3.27976173 | 1.56E-14 | 1.08E-12 | 2.87962109 | 2.28E-14 | 1.59E-12 |
| Il2 | -2.8195227 | | | 0.25147327 | | | -0.7021953 | | | -2.9179119 | | | -0.8913273 | | |
| Il33 | 0.36490963 | | | 0.44270045 | | | 0.44091491 | | | 0.22056838 | | | 0.23811717 | | |
| Il6 | 3.11479884 | 4.20E-05 | 0 | 2.00489386 | 0.006 | 0.01 | 1.32688326 | | | 1.91843524 | 0.01 | 0.07 | 0.55574405 | | |
| Irf4 | 0.09390505 | | | 0.96870481 | | | 1.08682157 | | | 1.56894436 | 0.01 | 0.08 | 1.51991042 | 0.03 | 0.19 |
| Irf5 | 1.34407122 | 0.01 | 0.04 | 1.97142312 | 0 | 0 | 0.96774616 | | | 1.61031704 | 0.005 | 0.05 | 1.7505996 | 0.005 | 0.05 |
| Myd88 | 1.14664026 | 0.01 | 0.04 | 1.30307144 | 0.005 | 0.01 | 0.43713789 | | | 0.91432487 | | | 0.72353769 | | |
| Nfkb1 | 0.27768821 | | | 0.5680915 | 0.03 | 0.07 | 0.23257477 | | | 0.42910188 | | | 0.25915407 | | |
| Nkg7 | 1.46460434 | | | 3.28311875 | 0 | 0 | 2.11834116 | 0.01 | 0.09 | 3.01591203 | 0.001 | 0.01 | 2.86095584 | 0.003 | 0.03 |
| Nos2 | 6.87477916 | 3.12E-13 | 8.68E-12 | 4.73589435 | 0.02 | 0.04 | 1.27919391 | | | 3.51951931 | 3.39E-09 | 1.18E-07 | 2.6596629 | | |
| Stat1 | 0.60692927 | | | 1.5003169 | 0 | 0 | 0.72283543 | | | 1.09022946 | 0.01 | 0.08 | 1.12299278 | 0.02 | 0.16 |
| Stat6 | 0.68416169 | | | 1.04042504 | 0.007 | 0.01 | 0.56874348 | | | 0.74609423 | | | 0.50843211 | | |
| Tnfrsf1a | 0.38892727 | | | 0.38531613 | | | -0.0588726 | | | -0.0593433 | | | -0.105608 | | |

**Table S2. Myogenesis related genes throughout the different time points after repeated simulated birth injuries.**

P and q values are indicated for significantly differentially expressed genes compared to uninjured group. FC= Log2FC

| | Days after 2nd Simulated Birth Injury | | | | | | | | | | | | | | |
|---|---|---|---|---|---|---|---|---|---|---|---|---|---|---|---|
| Gene | 3 | | | 7 | | | 21 | | | 35 | | | 63 | | |
| | FC | p | q | FC | p | q | FC | p | q | FC | p | q | FC | p | q |
| Pax7 | -0.09 | | | 0.16 | | | 0.37 | | | -0.25 | | | -0.26 | | |
| Acta1 | 0.02316936 | | | -0.3000812 | | | 0.58008995 | | | 0.56558299 | | | 0.46083841 | | |
| Actn2 | -0.3249137 | | | -0.0452203 | | | 0.17152183 | | | -0.1001479 | | | -0.1789316 | | |
| Adam12 | 2.25623937 | 0 | 0 | 2.41424305 | 0 | 0 | -0.2462 | | | -0.4756179 | | | -0.4449533 | | |
| Des | -0.3392639 | | | -0.3868541 | | | -0.1548618 | | | -0.2043479 | | | -0.1316071 | | |
| Dmd | -0.2629371 | | | -0.0392671 | | | 0.34949445 | | | 0.06310284 | | | 0.12853762 | | |
| Hgf | 0.58899434 | | | 0.76688986 | | | 0.14596382 | | | 0.46351249 | | | 0.19603451 | | |
| Mybpc1 | -0.9647268 | 0 | 0.02 | -0.9015688 | 0.01 | 0.02 | -0.4799291 | | | -0.593721 | | | -0.4656967 | | |
| Mybpc2 | -0.2110009 | | | -0.3021629 | | | 0.16940731 | | | 0.12526474 | | | 0.21432811 | | |
| Myf5 | 0.92194386 | 0 | 0 | 0.79706794 | 0 | 0 | 0.73801075 | 0.007 | 0.06 | 0.33679825 | | | -0.2852865 | | |
| Myf6 | -0.821629 | 0 | 0 | -0.5857951 | 0.009 | 0.02 | -0.8068009 | 0 | 0.005 | -0.6843907 | 0.003 | 0.03 | -0.3194646 | | |
| Myh1 | 0.01195634 | | | -0.2487938 | | | 0.52803377 | | | -0.0115645 | | | 0.18481157 | | |
| Myh3 | 2.81544654 | 0 | 0 | 5.25742584 | 2.71E-07 | 4.71E-06 | 2.65528892 | 0 | 0.03 | 0.75108399 | | | 0.27900562 | | |
| Myh7 | -0.317075 | | | -0.4144545 | | | 0.37514331 | | | 0.3214762 | | | 0.43807999 | | |
| Mylk2 | -0.5360198 | | | -0.9701489 | 0.04 | 0.08 | -0.2859813 | | | -0.2997333 | | | -0.1645212 | | |
| Myod1 | -0.7678297 | 0.01 | 0.04 | -1.2865482 | 4.87E-05 | 0 | -1.0082759 | 0 | 0.02 | -1.1671528 | 0 | 0.007 | -1.2080089 | 0.001 | 0.02 |
| Myog | 0.81063011 | 0 | 0.02 | 0.22940258 | | | 0.00566239 | | | -0.412943 | | | -0.506345 | | |
| Sln | 1.62742805 | 0.01 | 0.04 | 2.3211714 | 0 | 0 | 0.76373956 | | | 0.78811659 | | | -0.0686495 | | |
| Tcap | -0.2345825 | | | -0.806663 | 0.04 | 0.08 | -0.1391563 | | | -0.0671382 | | | 0.13954969 | | |
| Tmod1 | -0.7000945 | 0.03 | 0.07 | -0.9272155 | 0.004 | 0.01 | -0.6639865 | 0.04 | 0.19 | -0.4042975 | | | -0.4825921 | | |
| Tnnt1 | -0.1440658 | | | -0.2422662 | | | 0.35086716 | | | 0.35739569 | | | 0.4321472 | | |
| Tnnt3 | -1.0871174 | 0 | 0.02 | -1.0422206 | 0.009 | 0.02 | -0.4381052 | | | -0.6057715 | | | -0.5354041 | | |
| Tpm3 | -0.1672384 | | | -0.1447604 | | | 0.52567842 | | | 0.53401076 | | | 0.52503759 | | |
| Tpm4 | 1.11113252 | 0 | 0.02 | 1.1130846 | 0.005 | 0.01 | 0.56239046 | | | 0.72901099 | | | 0.44673377 | | |
| Ttn | -0.2838444 | | | -0.0821689 | | | 0.10928269 | | | 0.2159503 | | | 0.22653365 | | |

**Table S3. Muscle anabolism and catabolism related genes throughout the different time points after repeated simulated birth injuries.**

P and q values are indicated for significantly differentially expressed genes compared to uninjured group. FC= Log2FC

| | Days after 2nd Simulated Birth Injury | | | | | | | | | | | | | | |
|---|---|---|---|---|---|---|---|---|---|---|---|---|---|---|---|
| Gene | 3 | | | 7 | | | 21 | | | 35 | | | 63 | | |
| | FC | p | q | FC | p | q | FC | p | q | FC | p | q | FC | p | q |
| **Has1** | 1.63 | 0.02 | 0.05 | 1.08 | | | 0.81 | | | 1.24 | | | -2.49 | | |
| **Akt1** | -0.231723 | | | -0.1722273 | | | -0.2015771 | | | -0.229714 | | | -0.1886912 | | |
| **Akt2** | -0.4652612 | 0.03 | 0.07 | -0.4924408 | 0.02 | 0.04 | -0.2897174 | | | -0.3163823 | | | -0.2346297 | | |
| **Camkk1** | 0.34601646 | | | 0.67307267 | 0.01 | 0.03 | 0.31531649 | | | 0.20860987 | | | 0.0898435 | | |
| **Ctsl** | 0.66778891 | 0.03 | 0.08 | 0.53543334 | | | 0.13569182 | | | 0.14969831 | | | 0.17911889 | | |
| **Eif2b5** | -0.7510263 | 5.16E-05 | 0 | -0.9539257 | 2.32E-07 | 4.61E-06 | -0.5199577 | 0.005 | 0.05 | -0.5124645 | 0.008 | 0.05 | -0.5036815 | 0.02 | 0.1 |
| **Eif4b** | -0.239648 | | | -0.464939 | 0.006 | 0.01 | 0.0688716 | | | 0.00730212 | | | 0.08677879 | | |
| **Eif4e** | -0.6869186 | 0 | 0 | -0.4830864 | 0.009 | 0.02 | -0.4015353 | 0.03 | 0.15 | -0.4843876 | 0.01 | 0.07 | -0.3122299 | | |
| **Eif4ebp1** | 0.06499577 | | | -0.6718313 | 1.37E-08 | 3.19E-07 | -0.1354139 | | | -0.2080245 | | | -0.0157674 | | |
| **Eif4g1** | -0.3752678 | | | -0.5232425 | 0.005 | 0.01 | -0.1131203 | | | -0.2607579 | | | -0.1760917 | | |
| **Fbxo32** | 0.15592488 | | | -0.5352283 | | | 0.16298208 | | | 0.5890172 | | | 0.58180934 | | |
| **Foxo1** | -0.0684141 | | | 0.02047918 | | | 0.27164583 | | | 0.30360012 | | | 0.52989642 | 0.002 | 0.031 |
| **Foxo3** | 0.16472942 | | | 0.14308394 | | | 0.28564863 | | | 0.10835136 | | | 0.36890497 | 0.02 | 0.18 |
| **Fst** | -1.1383531 | 4.18E-05 | 0 | -0.4274297 | | | -1.0452085 | 0 | 0.004 | -0.6323033 | 0.03 | 0.1 | -1.5115167 | 8.31E-05 | 0.001 |
| **Grb10** | -0.7131634 | 4.52E-11 | 1.04E-09 | -0.4280018 | 6.61E-05 | 0 | -0.5581825 | 2.49E-07 | 6.90E-06 | -0.5826393 | 2.78E-07 | 5.52E-06 | -0.8550161 | 1.61E-10 | 5.61E-09 |
| **Gsk3b** | -0.3626213 | | | -0.1242114 | | | -0.0006434 | | | -0.0191362 | | | 0.04370119 | | |
| **Ikbkb** | -0.3944253 | | | -0.4117868 | | | -0.5497125 | 0.006 | 0.06 | -0.2987984 | | | -0.277723 | | |
| **Junb** | 2.10179447 | 5.39E-05 | 0 | 1.49297542 | | | 0.75966758 | | | 1.67224876 | | | 1.02101225 | | |
| **Mstn** | -0.9701743 | 0.03 | 0.08 | -1.8641435 | 5.50E-05 | 0 | -1.3631566 | 0.003 | 0.03 | -1.4342014 | 0.003 | 0.03 | -1.6709658 | 0.003 | 0.04 |
| **Mtor** | -0.1856479 | | | -0.4352534 | 0 | 0 | -0.0765345 | | | -0.0587486 | | | 0.07826 | | |
| **Nfkbia** | 0.48953087 | | | 0.34088626 | | | 0.12397139 | | | 0.51776502 | | | 0.55216476 | | |
| **Nol3** | 0.35462079 | | | -0.1948529 | | | 0.27075141 | | | 0.13782787 | | | 0.21164634 | | |
| **Prkaa1** | -0.1250985 | | | -0.0431805 | | | 0.16724791 | | | 0.06863348 | | | -0.1507732 | | |
| **Rheb** | -0.536009 | 0 | 0.01 | -0.5667091 | 0 | 0 | -0.2793008 | | | -0.2096313 | | | -0.1158169 | | |
| **Rps6ka1** | 0.84608062 | | | 1.45902051 | 0.004 | 0.01 | 0.48662641 | | | 1.18148563 | 0.02 | 0.1 | 1.07123279 | | |
| **Rps6kb1** | -0.7580955 | 0 | 0 | -0.4958105 | 0.01 | 0.03 | -0.4886425 | 0.01 | 0.09 | -0.5827838 | 0.005 | 0.05 | -0.6094505 | 0.01 | 0.1 |
| **Ccn4** | 1.66053332 | 0.01 | 0.04 | 1.66619993 | 0.01 | 0.03 | 0.28660856 | | | -1.3366488 | | | -1.4164371 | | |
| **Trim63** | -0.1194405 | | | -0.6658114 | 0.01 | 0.03 | -0.2713681 | | | -0.0601036 | | | 0.27408859 | | |

**Table S4. Extracellular matrix related genes throughout the different time points after repeated simulated birth injuries.**

P and q values are indicated for significantly differentially expressed genes compared to uninjured group. FC= Log2FC

| | Days after 2nd Simulated Birth Injury | | | | | | | | | | | | | | |
|---|---|---|---|---|---|---|---|---|---|---|---|---|---|---|---|
| Gene | 3 | | | 7 | | | 21 | | | 35 | | | 63 | | |
| | FC | p | q | FC | p | q | FC | p | q | FC | p | q | FC | p | q |
| **Mmp8** | 1 | | | 1.07 | | | -0.71 | | | 0.21 | | | -0.56 | | |
| **Col1a1** | 0.98372203 | | | 1.2010992 | | | -0.1228866 | | | -0.8810262 | | | -1.3146721 | | |
| **Col3a1** | 1.21037758 | 0.01 | 0.05 | 1.49166045 | 0.004 | 0.1 | 0.43752722 | | | -0.1674783 | | | -0.7081826 | | |
| **Itga1** | 0.04934163 | | | 0.59379572 | 0 | 0 | 0.33939788 | 0.03 | 0.15 | 0.43374693 | 0.009 | 0.05 | 0.26405894 | | |
| **Itga2** | 0.02620677 | | | 0.09542747 | | | 0.06254833 | | | -1.0862344 | 0.01 | 0.07 | -0.7405512 | | |
| **Itgb1** | 0.13898235 | | | 0.2707636 | | | 0.03145044 | | | -0.0265108 | | | -0.1522956 | | |
| **Mmp1** | 0.45294773 | | | 0.17730176 | | | -0.0517942 | | | 0.09056389 | | | -1.709139 | | |
| **Mmp2** | 0.27521354 | | | 1.09396247 | 0.008 | 0.02 | 0.49178323 | | | 0.24604027 | | | -0.0758155 | | |
| **Mmp9** | 1.08885893 | | | 1.65737323 | 0.01 | 0.03 | 0.393747 | | | -0.1329704 | | | -0.7321961 | | |
| **Pdgfa** | -0.6163226 | | | -0.3086441 | | | -0.3578908 | | | -0.2329094 | | | -0.8608714 | 5.00E-08 | 1.41E-06 |
| **Pdfgra** | 0.31025163 | | | 0.41523157 | | | -0.0863701 | | | 0.08351219 | | | -0.4507678 | | |
| **Smad3** | -0.9829924 | 0 | 0 | -1.0431873 | 0 | 0 | -0.7334231 | 0.01 | 0.08 | -0.7788297 | 0.009 | 0.05 | -0.9574591 | 0.005 | 0.05 |
| **Smad4** | -0.2889801 | | | -0.2002076 | | | -0.0571093 | | | -0.0338918 | | | 0.18623242 | | |
| **Tgfb1** | 1.14273908 | 0.03 | 0.08 | 1.76862926 | 0 | 0 | 0.83367159 | | | 1.19081527 | 0.03 | 0.12 | 1.08073156 | | |
| **Tgfbr1** | 0.19425668 | | | 0.22106398 | | | 0.11303336 | | | 0.22301536 | | | 0.20326674 | | |
| **Timp1** | 2.84303327 | 3.95E-05 | 0 | 2.21586164 | 0 | 0 | 0.72818828 | | | 1.27942783 | 0.04 | 0.1 | 0.65999542 | | |
| **Timp2** | -0.1740448 | | | 0.36634178 | | | 0.23366219 | | | 0.19095401 | | | -0.0442825 | | |
| **Timp3** | -0.2785393 | | | -0.2382331 | | | 0.49908612 | | | 0.2056212 | | | 0.35892781 | | |
| **Timp4** | -0.3162146 | | | -0.2132306 | | | 0.15182609 | | | 0.1179532 | | | 0.44529833 | | |
| **Vwa1** | 0.61124212 | | | 0.99136444 | 0.01 | 0.03 | 0.18669157 | | | 0.20776831 | | | 0.01139593 | | |

**Table S5. Neuromuscular junctions related genes throughout the different time points after repeated simulated birth injuries.**

P and q values are indicated for significantly differentially expressed genes compared to uninjured group. FC= Log2FC

| | Days after 2nd Simulated Birth Injury | | | | | | | | | | | | | | |
|---|---|---|---|---|---|---|---|---|---|---|---|---|---|---|---|
| Gene | 3 | | | 7 | | | 21 | | | 35 | | | 63 | | |
| | FC | p | q | FC | p | q | FC | p | q | FC | p | q | FC | p | q |
| Dvl1 | -0.85 | 8.92E-06 | 0 | -0.58 | 0 | 0.01 | -0.51 | 0.007 | 0.06 | -0.53 | 0.007 | 0.05 | -0.51 | 0.02 | 0.1 |
| Agrn | 0.07102083 | | | 0.42878044 | | | 0.03684727 | | | -0.2860692 | | | -0.265273 | | |
| App | 0.1228353 | | | 0.47255595 | 0.03 | 0.06 | 0.02390995 | | | 0.16446805 | | | -0.0030643 | | |
| Erbb2 | 1.04988573 | 0 | 0.03 | 0.78169282 | 0.04 | 0.08 | 0.41891115 | | | 0.28719009 | | | 0.53654034 | | |
| Gphn | -0.397876 | | | -0.4483913 | | | -0.1131154 | | | -0.2095773 | | | -0.0122485 | | |
| Musk | 0.79191941 | | | 1.13027991 | 0.01 | 0.03 | -0.0462679 | | | -0.0948886 | | | -0.2371049 | | |
| Nrg1 | 1.24181907 | 0 | 0.01 | 1.51925473 | 8.40E-05 | 0 | 0.1231052 | | | 1.12088398 | 0.02 | 0.1 | 0.811348 | | |
| Ptn | 1.14273933 | 0.01 | 0.04 | 1.16021009 | 0.01 | 0.03 | 0.36964083 | | | 0.52350832 | | | -0.16198 | | |
| Tnc | 2.33961511 | 0 | 0.01 | 2.42767138 | 0 | 0.01 | 0.00639764 | | | -0.0570216 | | | -0.9897771 | | |
| Utrn | -0.3686152 | | | -0.0802778 | | | -0.1811844 | | | -0.0649315 | | | -0.2013801 | | |

**Table S6. Vascularization related genes throughout the different time points after repeated simulated birth injuries.**

P and q values are indicated for significantly differentially expressed genes compared to uninjured group. FC= Log2FC

| | Days after 2nd Simulated Birth Injury | | | | | | | | | | | | | | |
|---|---|---|---|---|---|---|---|---|---|---|---|---|---|---|---|
| Gene | 3 | | | 7 | | | 21 | | | 35 | | | 63 | | |
| | FC | p | q | FC | p | q | FC | p | q | FC | p | q | FC | p | q |
| Thsb1 | 1.7 | 0 | 0.02 | 1.81 | 0.004 | 0.01 | 0.5 | | | 0.57 | | | 0.03 | | |
| Angpt1 | -0.562359 | 0.04 | 0.09 | -0.4918092 | | | 0.44142617 | | | -0.1530769 | | | 0.14926645 | | |
| Angpt2 | -0.0535874 | | | 0.48069606 | 0.01 | 0.03 | 0.27448998 | | | 0.44452322 | 0.03 | 0.1 | -0.1156 | | |
| Fgf2 | -1.4901025 | 0.0001 | 0.001 | -0.8692082 | 0.02 | 0.04 | -1.0405345 | 0.007 | 0.06 | -0.943594 | 0.02 | 0.1 | -0.8296596 | | |
| Flt1 | -0.0411508 | | | -0.1269556 | | | -0.0514609 | | | 0.02596042 | | | -0.0590792 | | |
| Kdr | -0.8413409 | 0 | 0 | 0.16400614 | | | -0.4384014 | | | -0.6412035 | 0.02 | 0.1 | -0.6169142 | | |
| Mmp14 | 0.65281754 | | | 1.58154279 | 0.002 | 0.01 | 0.47101963 | | | 0.63130721 | | | -0.1493721 | | |
| Nrp1 | -0.1903515 | | | 0.37244373 | | | -0.0837235 | | | -0.0273247 | | | -0.2683051 | | |
| Tie1 | 0.27733478 | | | 0.4818113 | | | 0.23309979 | | | 0.19527437 | | | 0.11773356 | | |
| Vegfa | -1.0128641 | 0 | 0 | -0.8615957 | 0 | 0 | -0.6542518 | 0.01 | 0.09 | -0.6074338 | 0.03 | 0.1 | -0.7111762 | 0.03 | 0.18 |
| Vegfb | -0.9132063 | 0 | 0 | -0.6642029 | 0.006 | 0.01 | -0.4725045 | | | -0.6661703 | 0.008 | 0.05 | -0.4278072 | | |
